# Canonical nuclear envelope protein emerin regulates structure and integrity of the erythrocyte plasma membrane

**DOI:** 10.1101/2025.06.06.658339

**Authors:** Brendan J. O’Brien, Joyleen Oliver, Evrett Thompson, Madeline Y. Mayday, Rubia Mancuso, Alokkhumar Jha, Jennifer Kwan, Gisèle Bonne, Rabah Ben Yaou, France Leturcq, Suxia Bai, Tim Nottoli, John Hwa, Diane S. Krause, Patrick G. Gallagher, Sean X. Gu, Kathleen A. Martin

## Abstract

Red blood cells (RBC) are essential for the survival of aerobic organisms. Mutations in RBC plasma membrane proteins can give rise to a spectrum of diseases characterized by hemolysis, inefficient gas exchange, and anemia. We tested whether emerin (EMD), a nuclear envelope (NE) protein, is also expressed in anuclear cell types to regulate membrane structure and function. We find that emerin is localized to the RBC plasma membrane and regulates the localization of other membrane proteins involved in actin branching and cytoskeletal dynamics. Additionally, we find that loss of *Emd* in mice results in decreased RBC corpuscular volume and hemoglobin content in both sexes, as well as a male-specific increase in RBC number and hematocrit, consistent with other male-dominated *EMD* phenotypes. Along with decreased RBC size, we observed a reduction in the rate of osmotic lysis in *Emd* KO animals, particularly males. We found that human *EMD* mutations also have a statistically significant association with RBC phenotypes, namely MCV. Individuals with emerin-related Emery-Dreifuss muscular dystrophy (EDMD) also demonstrate a mild yet consistent RBC phenotype according to patient CBC data. Together, these data suggest that emerin can have cell-specific localization and functions independent of its canonical repertoire in nucleated cells.

## Introduction

Erythrocytes (RBCs) play a vital role in human physiology, primarily by transporting oxygen from the lungs to peripheral tissues and retrograde transport of carbon dioxide for exhalation^1^. As such, dysfunction in RBC physiology can lead to various disorders affecting oxygen^2,3^. Mutations in genes encoding RBC membrane proteins can have dramatic effects on cell shape, flexibility, rate of gas exchange, and sensitivity to hemolysis. RBCs normally have a biconcave disc shape that optimizes surface area for gas exchange and allows for deformation as these cells traverse capillaries. The most common RBC membrane disorder is hereditary spherocytosis, which is a condition marked by spherical-shaped RBCs due to a deficiency in proteins including spectrin^4^, ankyrin^5^, or band 3^6-8^. The altered shape makes the RBCs more fragile and susceptible to premature rupture, resulting in anemia, jaundice, and splenomegaly. Other RBC membrane disorders include elliptocytosis, pyropoikilocytosis, microcytosis and stomatocytosis, among others^9,10^. The mechanisms regulating many of these diseases are an active area of investigation and therapeutic development.

Emerin (Emd) is a ubiquitously expressed 34 kDa protein localized in the inner nuclear membrane and has reported roles regulating nuclear shape, gene expression, cell signaling, and chromatin architecture in multiple cell types^11-14^. Mutations in the human emerin gene, *EMD*, contribute to a specific envelopathy referred to as Emery-Dreifuss muscular dystrophy (EDMD), inherited in an X-linked fashion affecting males, characterized by skeletal muscle wasting, joint contracture of major tendons and spine, and cardiac conduction and rhythm defects^15^. Female carriers are usually asymptomatic or manifest mild isolated cardiac disease as they carry normal and mutated alleles on their X chromosomes. Despite its well-accepted role in the inner nuclear membrane, emerin has also been reported to localize to other subcellular compartments such as the plasma membrane^16^, outer nuclear membrane and the endoplasmic reticulum following mechanical stimulation or viral infection^17-20^. In this study, we tested whether emerin is expressed in anuclear mature erythrocytes and whether extra-nuclear localization confers cell type-specific functions in these cells. Using genetic mouse models, bioinformatics, and human patient data, we found that emerin is indeed expressed in mature RBCs and regulates RBC size and membrane stability. These data suggest that emerin has cell-specific localization and functions beyond those reported by decades of work focused on the nuclear envelope.

## Methods

### Generation of *Emd* KO mouse

The constitutively deletion of emerin (*Emd* KO) mouse model was generated by first generating a mouse line with LoxP sites flanking exons 2-4 of the *Emd* gene, then crossing this mouse with an *Actb*-Cre to permanently excise the *Emd* sequence in all cells. The floxed mouse was generated via CRISPR-Cas9 genome editing^21-23^. Potential Cas9 target guide (protospacer) sequences in introns 1, 4, and 5 were screened using the online tool CRISPOR http://crispor.tefor.net^24^ and candidates were selected. Templates for sgRNA synthesis were generated by PCR, sgRNAs were transcribed *in vitro* and purified (Megashortscript, MegaClear; ThermoFisher). sgRNA/Cas9 RNPs were complexed and tested for activity by zygote electroporation, incubation of embryos to blastocyst stage, and genotype scoring of indel creation at the target sites. The sgRNAs that demonstrated the highest activity and would enable a single-stranded DNA recombination template to target both loxP sites simultaneously, were selected for creating the floxed allele. Guide RNA (gRNA) sequences are as follows: intron 1, 5’ guide: GTGGCGCTCAGAACCTCTCT and intron 4, 3’ guide GTTGGCTGGTAGGTATACCT. Guide primers for generating the template for transcription included a 5’ T7 promoter and a 3’ sgRNA scaffold sequence and were as follows (protospacer sequence underlined): 5’*TGTAATACGACTCACTATAGG*<u>GTGGCGCTCAGAACCTCTCT</u>*GTTTTAGAGCTAGAAATAGC* 3’ 5’*TGTAATACGACTCACTATAGG*<u>GTTGGCTGGTAGGTATACCT</u>*GTTTTAGAGCTAGAAATAGC* 3’ sgRNA template reverse primer: 5’ AAAAGCACCGACTCGGTGCC 3’ Accordingly, a 1206 base long single-stranded DNA (lssDNA) recombination template incorporating the 5’ and 3’ loxP sites at the Cas9 cleavage sites was synthesized (IDT). The injection mix of sgRNA/Cas9 RNP + lssDNA was microinjected into the pronuclei of C57Bl/6J zygotes^23^. Embryos were transferred to the oviducts of pseudopregnant CD-1 foster females using standard techniques^25^. Genotype screening of tissue biopsies from founder pups was performed by PCR amplification and Sanger sequencing to verify the floxed allele. Germline transmission of the correctly targeted allele (i.e., both loxP sites in cis) was confirmed by breeding and sequence analysis.

### Membrane Ghost Isolation

Peripheral blood from *Emd* KO and wild type (WT) mice was collected *via* cardiac puncture into citrate tubes to prevent coagulation. RBCs were spun down for one minute at 1,000 x g to remove the WBC and plasma layer. Cells were then resuspended and washed in ice-cold PBS containing protease inhibitors (Thermo Scientific, cat. #78447) followed by ice-cold PBS and centrifuged at 2500 RPM for 10 minutes at 4°C. Cell pellet was then resuspended with Tris-EDTA (pH 8.0) lysis buffer and allowed to incubate on ice for 15 minutes. Cell suspension was then centrifuged for 10 minutes at 16,000 RPM at 4°C. The supernatant was removed and kept as the soluble fraction. RBC pellets were continuously resuspended in lysis buffer and centrifuged until the cell pellet was fully white in color, lacking all red hue. After the final spin, RBC ghosts were resuspended in sample buffer and boiled at 95°C for 5 minutes, spun, and used for western blot analysis. Additional, large aliquots of ghost fractions were kept at -80°C for long term storage.

### Western Blot

Western blots were performed as described previously in Martin lab publications^26,27^. Briefly, 10-20μg of proteins was loaded in each lane of 4-15% SDS-PAGE gels, transferred to nitrocellulose membranes (Biorad, P1620115), blocked with 5% nonfat dry milk, washed in Tris-buffered saline, 0.1% Tween 20 (TBS-T) and probed with polyclonal anti-Emerin antibody (Thermo Scientific, PA5-51424) and/or anti-ACTB (Sigma, A2228) overnight at 4°C. Membranes were incubated with HRP conjugated secondary antibodies (Thermo Scientific, 31460) washed in TBS-T, developed with Super signal West Pico Maximum Sensitivity Substrate (Pierce) and analyzed with the G:BOX imaging system (Syngene).

### Sample preparation for proteomics analysis

RIPA buffer (200 µL) (Thermo Scientific, cat. #89901) containing 1× HALT protease/phosphatase inhibitor cocktail (Thermo Scientific, cat. #78447) was added to each sample. The samples were vortexed for 3 min then centrifuged at 4 °C for 10 min at 20,500 g. A 150 µL aliquot of supernatant was removed from each sample and reduced with 4 mM dithiothreitol (Thermo Scientific, cat. #20290) for 30 min at 37 °C. Samples were cooled then alkylated with 8 mM iodoacetamide (Sigma-Aldrich, cat. #I1149) for 30 min at room temperature in the dark. Proteins were precipitated using chloroform/methanol/water and protein pellets were resuspended in 2 M urea/111 mM ammonium bicarbonate with water bath sonication. Proteins were digested at 37 °C via overnight incubation with 4 µg endoproteinase LysC (New England Biolabs, cat. #P8109S) followed by a 6.5 h incubation with 4 µg trypsin (Promega, cat. #V5113). Trifluoroacetic acid (TFA) was added to each sample to a final concentration of 1% and samples were desalted using BioPureSPN Macro columns (The Nest Group, Inc., cat. #HMM-S18V). Desalted peptides were dried via vacuum centrifugation and resuspended in 2% acetonitrile/0.2% TFA in water. Peptide concentrations were measured via absorbance at 280 nm using a Nanodrop spectrophotometer and 0.08 µg/µL dilutions were prepared. Pierce retention time calibration mixture (Thermo Scientific, cat. #88321) (12.5 fmol/µL) was added to each sample prior to injection.

### LC-MS/MS data acquisition

LC-MS/MS data were acquired on a Thermo Scientific Orbitrap Fusion mass spectrometer coupled to a Waters nanoACQUITY UPLC system. Sample injection order was randomized, and 400 ng of peptide were injected per sample. Peptides were loaded onto a Symmetry C18 trap column (100 Å, 5 µm, 180 μm × 20 mm) (Waters, cat. #186007496) at a flow rate of 5 µL/min and separated using a peptide BEH C18 analytical column (130 Å, 1.7 μm, 75 μm × 250 mm) (Waters, cat. #186007484) (45 °C). The compositions of mobile phases A and B were 0.1% formic acid in water and 0.1% formic acid in acetonitrile, respectively. The peptides were eluted at a flow rate of 300 nL/min with a gradient starting at 3% B, increasing to 5% B over 5 min, then to 20% B over 120 min and 35% B over 45 min. The gradient was then ramped to 97% B over 5 min and held for 5 min before the column was equilibrated to starting conditions. Precursor MS1 scans (profile) were collected in the orbitrap from 350-1550 m/z at a resolution of 120,000. The AGC target was set to 4 × 105 and the maximum injection time was 60 ms. Data-dependent MS2 scans (centroid) were collected in the ion trap starting at 100 m/z using the turbo scan rate setting. Precursors with 2 ≤ z ≤ 8 were selected for HCD fragmentation with a collision energy setting of 28%. The AGC target was set to 5 × 104 and the maximum injection time was 50 ms. The cycle time was set to 3 s and dynamic exclusion was enabled with a repeat count of one and an exclusion duration of 30 s. Four blank injections were conducted in between sample injections to minimize carryover.

### Proteomics data analysis

Protein identification and label-free quantification (LFQ) were performed in Proteome Discoverer (Thermo Fisher Scientific, version 3.0.1.27) using standard processing and consensus workflows. Data were searched against the SwissProt Mus musculus database (downloaded March 2024) and a database of common contaminant proteins using the Sequest HT search engine. Trypsin was selected as the enzyme, up to two missed cleavages were allowed, and precursor and product ion mass tolerances were set to 10 ppm and 0.6 Da, respectively. Oxidation of methionine was set as a variable modification and carbamidomethylation of cysteine was set as a fixed modification. Intensity-based rescoring of peptide spectral matches was performed using INFERYS prior to false discovery rate (FDR) estimation using Percolator. Peptide- and protein-level FDRs were set to 1%. Quantification was performed using unique peptides and normalization between files was based on total peptide amount. Proteins marked as master proteins with at least two unique peptides were exported for analysis.

### Fluorescent Labeling/Confocal Microscopy

Freshly isolated peripheral blood was collected from WT or *Emd* KO mice via cardiac puncture. Samples were collected directly into citrate tubes to inhibit coagulation. Samples were briefly spun down for one minute at 1,000 x g and resuspended in Dulbecco’s PBS. Following a second spin, 20 μL of RBC pellet was added to 50 μL of 3% volume glutaraldehyde in PBS. The fixation mix was allowed to incubate at room temperature for 15 minutes before being washed three times with 500 μL PBS. RBC pellets were then blocked and permeabilized with 0.1% Triton-X (Thermo Scientific cat. #28314) detergent and 2.5% normal goat serum (Cell Signaling Technologies cat. #5425S) in PBS for 30 minutes at room temperature. Following three additional washes, RBCs are incubated with antibody against emerin (Thermo Scientific cat. #PA5-51424) overnight at a concentration of 1:200 by volume. Additional washes proceed the addition of fluorescent secondary antibody and AlexaFluor568-conjugated phalloidin (Invitrogen cat. #A12380) stain, both at a concentration of 1:400 by volume. After a final set of washes, RBC pellets were resuspended in a 50% glycerol solution in PBS and mounted onto microscopy slides. Labeled RBCs were then imaged using a Leica sp8 inverted confocal microscope with a 63x oil objective. Z-stack image series were collected at 512 × 512 and exported as .lif files.

### Surface Rendering

Images series collected from the confocal microscope were brought to the Imaging Core at the Yale School of Medicine. There, Imaris 3D object-based rendering software version 10.0.1 (Oxford instruments) was used for image analysis of multiple confocal z-stack series to visualize RBC membrane shape. The surface module was utilized to render the RBC plasma membrane based off fluorescent intensity of phalloidin staining. The dot render creation wizard was used to visualize Emerin localization within the cells and color code loci based off proximity to the rendered surface. Representative images were exported with uniform contrast and brightness.

### Complete Blood Count

Blood was collected by cardiac puncture into a syringe with a ratio of 1:9 sodium citrate/blood and sent immediately to the laboratory. Complete blood count (CBC) was performed on the XN-9000 automated hematology analyzer (Sysmex, Inc.).

### Osmotic Fragility Test

Blood was collected via cardiac puncture as previously described. Beforehand, buffered 10% NaCl stock solution was made and stored at -20°C. Frozen stocks were used to make serial dilutions of 0.0, 0.55, 0.60, 0.65, and 0.85% saline. Blood (20 μL) was added to a microfuge tube of each saline solution and incubated at room temperature for 15 minutes. Samples were then centrifuged at 3000 rpm for 10 minutes. Supernatants were added to a 96 well format ELISA plate and absorbances were read using a BioTek SYNERGY H1 microplate reader. Percent hemolysis was calculated by the following equation:

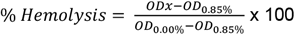

### Bioinformatic Analysis of *EMD* variants

Data were sourced from the ExPheWAS platform^28^ and the Pan-UK Biobank^29^ to ensure comprehensive phenotype coverage. Genotypic and phenotypic associations were extracted from the ExPheWAS platform, which provided insights into cardiometabolic, immune, and other disease-related phenotypes. Additional phenotype data were obtained from the Pan-UK Biobank, offering a rich dataset of genomic and health-related traits. Variant annotation was performed using the GenoHub platform^30^ to identify genomic variants linked to the *EMD* gene. Cell-type-specific analysis of these variants was conducted using PCGA tools^29^ to assess variant enrichment in specific cell populations. To integrate phenotype data from the ExPheWAS and Pan-UK Biobank, we employed Deep Phenotype-Variant Prioritization (Deep PVP)^31^, a machine learning model designed to harmonize and prioritize phenotypic associations. Deep PVP utilized key features, including statistical significance, effect size, phenotype frequency, and cross-cohort consistency, to evaluate and rank phenotypes associated with each variant. The model was trained on curated datasets to learn patterns of robust associations and mitigate biases arising from platform-specific data. Deep PVP employed a neural network architecture to account for complex, non-linear relationships between features and incorporated attention mechanisms to identify the most predictive signals. The model provided a confidence score for each phenotype-variant pair, emphasizing phenotypes with consistent and significant associations across datasets. Missing data and platform inconsistencies were addressed using imputation techniques and ensemble averaging. The integration and prioritization enabled by Deep PVP yielded a ranked list of phenotype-variant associations, highlighting biologically meaningful relationships with high confidence. This method facilitated the identification of the best possible phenotypes linked to *EMD* variants, providing a foundation for downstream analysis and interpretation.

## Results

### Emerin is expressed in mature RBCs and enriched in isolated membrane fractions

To evaluate whether emerin is expressed in non-nucleated, mature RBCs, we isolated whole blood *via* cardiac puncture from wild-type (WT) and *Emd* knockout (*Emd* KO) mice. We used a validated method to enrich the membrane fraction of the RBCs via isotonic lysis to create membrane “ghosts”, removing cytosolic components such as hemoglobin^32^. After enrichment of plasma membranes following isotonic lysis, emerin protein expression was readily detectable in murine RBC membrane ghosts (**Fig. 1A**). Additionally, we validated our novel *Emd* KO strain (see Methods), which demonstrated loss of emerin expression in the RBC membrane under the same experimental conditions in both sexes (**Fig. 1A, 1B**). To assess the emerin-dependent proteomic landscape of the RBC membrane, we performed mass spectrometry on membrane ghosts isolated from WT and *Emd* KO mice. We were unable to identify any emerin peptides in our *Emd* KO group, providing secondary validation of complete knockout (**Fig. 1C**). We observed robust separation between WT and KO samples in principle component analysis (PCA) (**Fig. 1D, left**) and identified that *Emd* KO resulted in hundreds of differentially expressed proteins within the membrane fraction when compared to WT counterparts (**Fig. 1E**). We utilized Qlucore enrichment analysis software to calculate proteins with at least 60% association with loss of emerin expression with FDR < 0.05 when comparing the two groups and performed pathway enrichment analyses (**Fig. 1D, right**). In the protein-protein interaction database (PPI-CORUM) documenting experimentally verified interactions, proteins downregulated by *Emd* deletion showed significant enrichment in cytoskeletal protein complexes related to actin branching protein ARP2/3, cofilin/CAP, and the septin family of proteins, among others. As a positive control, we also identified significant enrichment of proteins in “Emerin complex 52”, including known emerin-binding proteins like Lmnb1 and Iqgap1^33^. Notably, proteins that were differentially expressed in the absence of emerin include several known to play roles in aberrant RBC physiology, like Band3, Band4, and Stom^6,9^. Additionally, Mammalian Phenotypic analysis (Monarch) supported that RBC parameters like mean corpuscular volume (MCV) and mean corpuscular hemoglobin (MCH) could potentially be affected by differential protein expression in *Emd* KO cells. (**Fig. 1F**). Together, these data suggest that emerin is enriched in the plasma membrane of mature RBCs and may play a key role in membrane structure and integrity.

**Figure 1.**
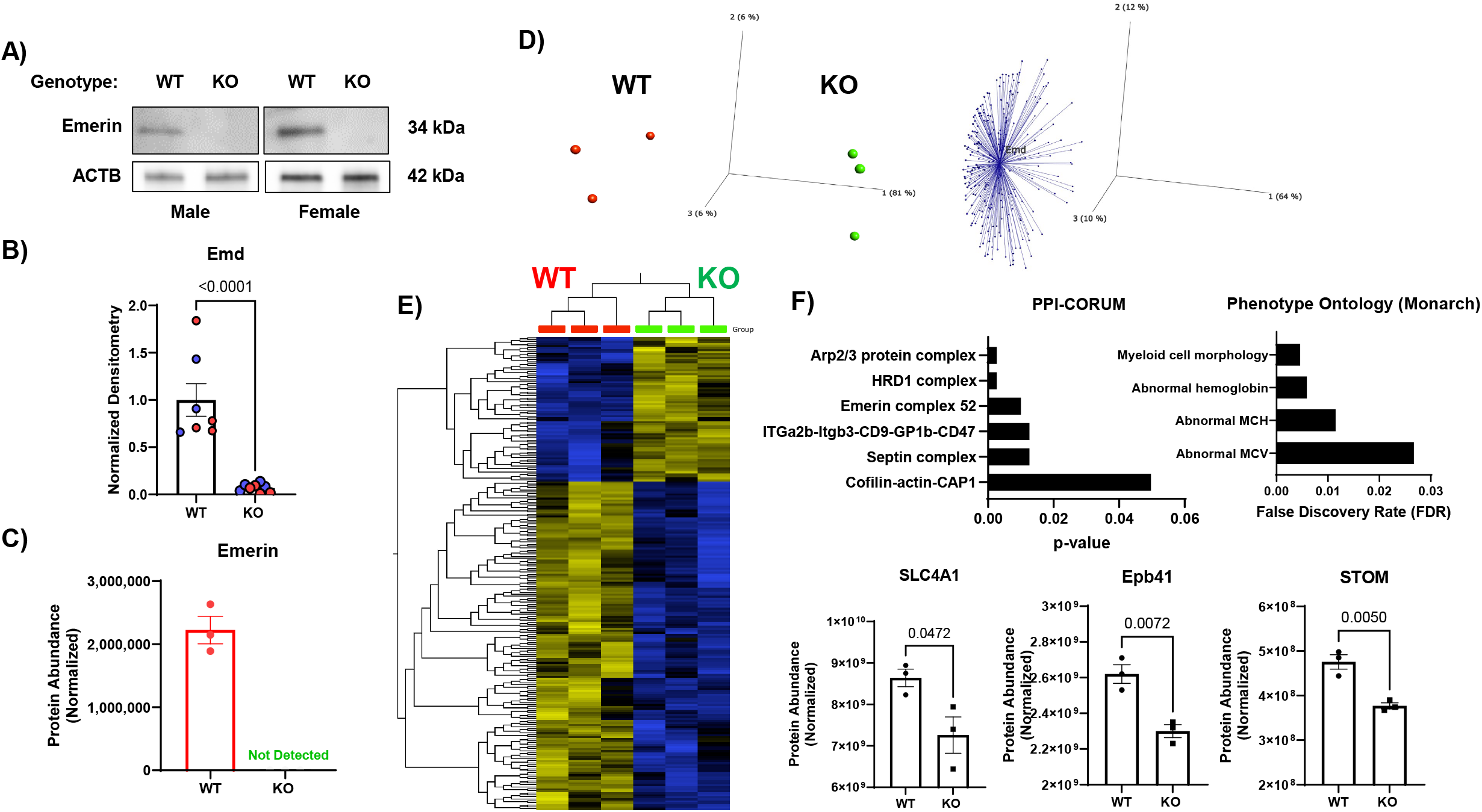
Emerin is expressed in mature RBCs and is enriched in isolated membrane fractions. **A)** Western blotting of emerin in murine RBC membrane ghosts isolated from WT and homozygous KO male and female mice. **B)** Quantitation of emerin expression normalized to Actb densitometry as analyzed with ImageJ. n = 3 of each sex of each respective genotype. Data represented as Mean ± SEM, Student’s t test. **C)** Normalized protein abundances of emerin peptides identified by mass spec proteomic analysis of isolated RBC membrane ghosts. n = 3 for each genotype. No signal was detected for emerin in *Emd* KO group. **D)** Principal component analysis of the six submitted ghost membrane samples (left) and correlation analysis (Qlucore, right) showing identified proteins with 60% correlation of expression with emerin expression. **E)** Heatmap generated in Qlucore of differentially expressed proteins between *Emd* KO ghost membranes (green) and WT counterparts (red). **F)** Protein-protein interaction CORUM and Mammalian Phenotype Ontology enrichment of proteins associated with emerin expression (60% and FDR<0.05). Select proteins represented as Mean ± SEM, Student’s t-test.

### Emerin is a critical regulator of RBC morphology and composition

To test more directly if emerin regulates RBC membrane morphology and intracellular content, we performed complete blood count (CBC) of washed RBCs from male and female mice from both WT and homozygous *Emd* KO strains. *Emd* knockout resulted in a significant decrease in mean corpuscular volume (MCV) and hemoglobin (MCH) in both sexes without a change in mean corpuscular hemoglobin concentration (MCHC) compared to WT counterparts (**Fig. 2A-D**). These data also revealed a compelling sex-specific phenotype: in male mice, loss of emerin expression resulted in an increase in RBC number (**Fig. 2E**), hemoglobin (**Fig 2F**), and hematocrit (**Fig 2G**), whereas female mice showed statistically insignificant changes, but with consistent trends. In humans, it is well-reported that mutations in *EMD* give rise to male-dominated phenotypes since *EMD* is located on the X-chromosome^34-36^. Given that both sexes of our knockout animals are fully deficient in emerin expression (**Fig. 1B**), sex itself appears to be a major contributing variable to emerin-dependent RBC phenotypes independent of protein expression. We did not observe a statistically significant alterations in the counts of the other circulating cell types, including platelets, lymphocytes, monocytes, or nucleated RBCs. We did, however observe a significant decrease in the number of circulating neutrophils in male *Emd* KO mice compared to WT counterparts (**Fig. 3**). These data are particularly compelling given the unique morphology and function of the nucleus in neutrophil biology^37^ and are the subject of a parallel investigation.

**Figure 2.**
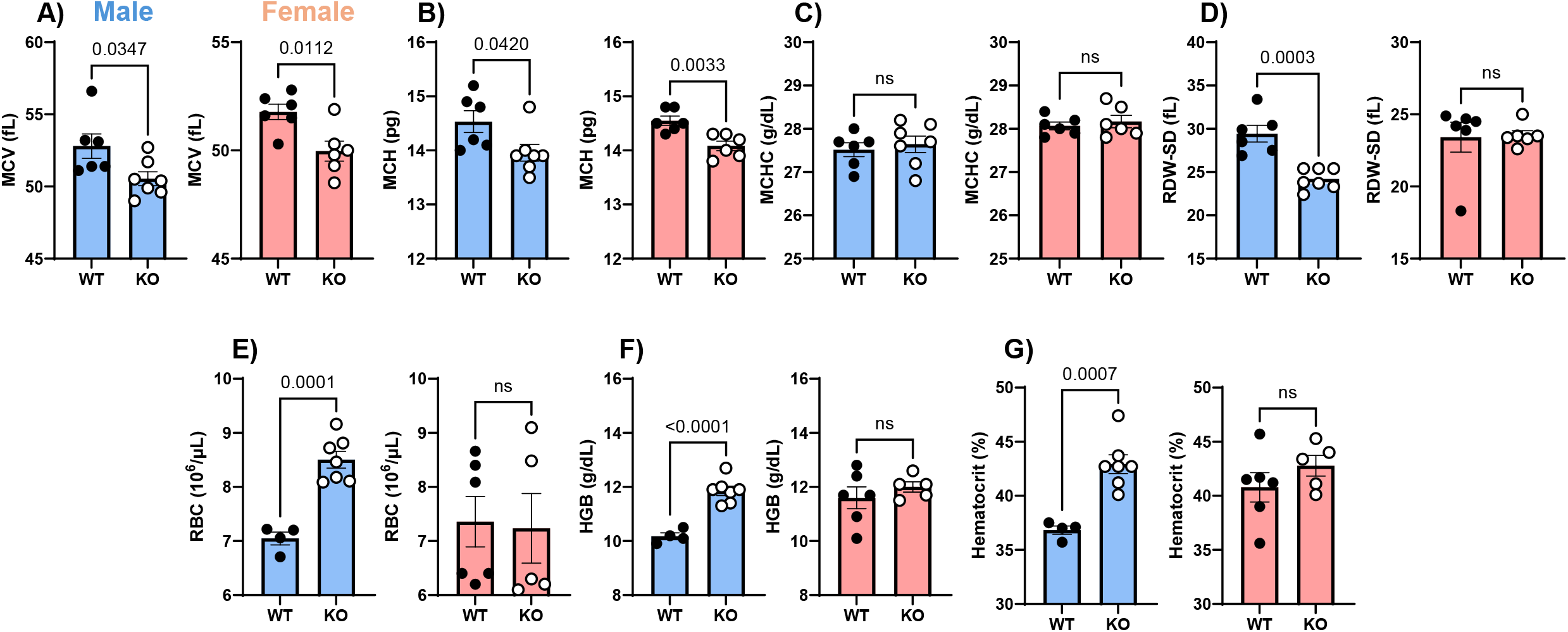
Effect of *Emd* knockout on RBC parameters *in vivo*. Complete blood counts (CBC) of freshly isolated peripheral blood from male (left) and female (right) mice: **A)** MCV, **B)** MCH, **C)** MCHC, **D)** RDW-SD, **E)** RBC number, **F)** hemoglobin, and **G)** hematocrit. Male mice shown as blue bars, female mice shown as red bars. n = 6 WT males, 7 KO males, 6 WT females, and 6 KO females. Data represented as Mean ± SEM. Student’s t-test.

**Figure 3.**
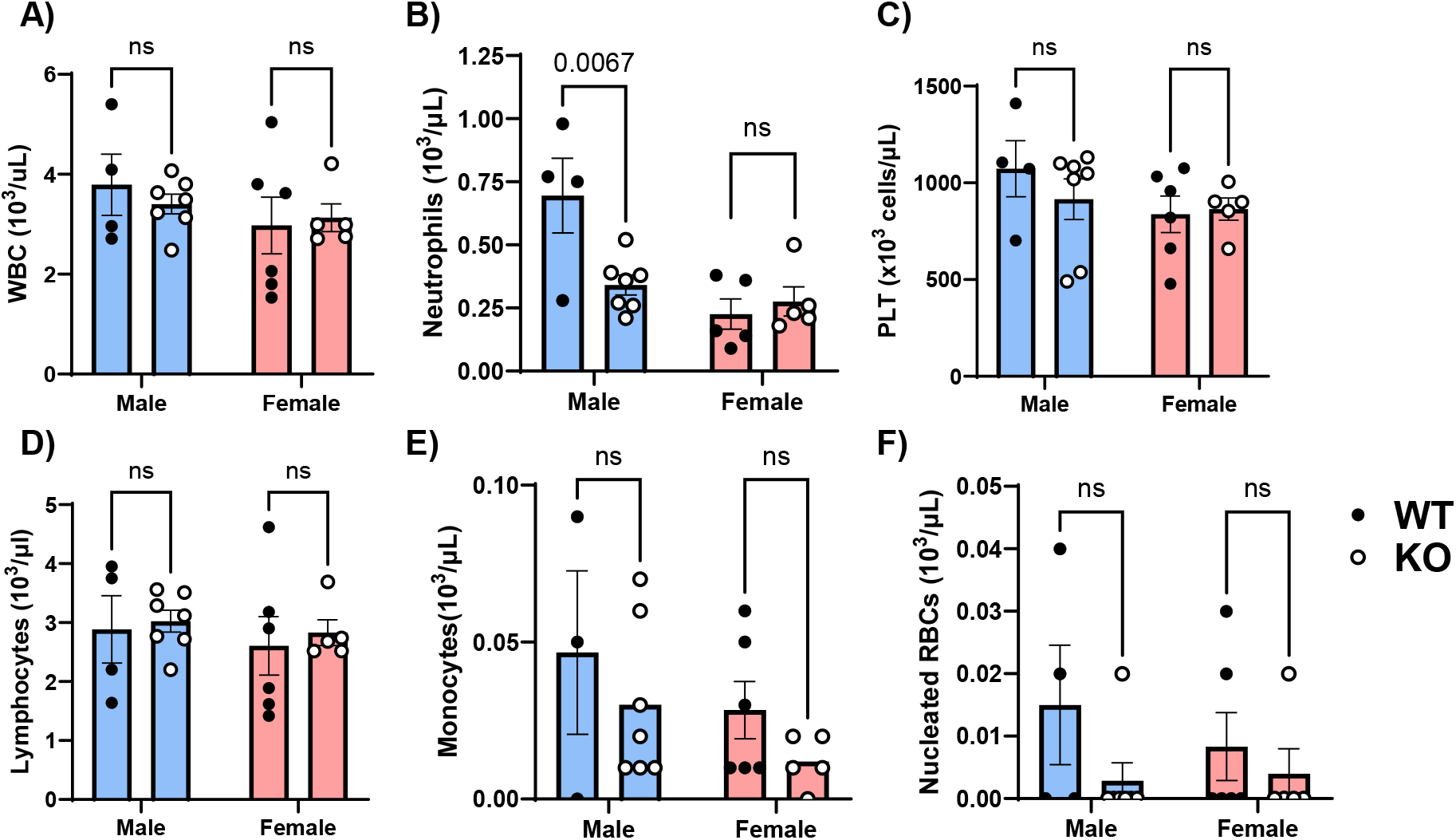
Effect of *Emd* knockout on other circulating cell types. Complete blood counts (CBC) of freshly isolated peripheral blood from male (left) and female (right) mice: **A)** white blood cells, **B)** Neutrophils, **C)** platelets, **D)** lymphocytes, **E)** monocytes, and **F)** nucleated RBCs. Male mice shown as blue bars, female mice shown as red bars. n = 3-5 WT males, 7 KO males, 6 WT females, and 6 KO females. Data represented as Mean ± SEM. Student’s t-test.

To further investigate the RBC phenotypes observed in *Emd* KO mice, we utilized high magnification fluorescence microscopy to directly visualize RBC structure. We validated our earlier data that emerin is expressed in mature WT RBCs (**Fig 1A**) by immunofluorescent visualization of emerin and colocalization with phalloidin-labeled actin (**Fig. 4A, top**). We also provided additional validation of *Emd* knockout with no fluorescent signal observed in *Emd* KO mice at uniform laser intensity and gain (**Fig. 4A, bottom**). We utilized the Imaris imaging processing software to perform spot and surface rendering analyses based on confocal z-stack series to show proximity of emerin and phalloidin+ actin staining and to quantify spatial parameters of RBCs from each respective genotype (**Fig. 4B**). Using the quantitative tools in the surface rendering module, we observed that both male and female homozygous *Emd* KO mice have decreased surface area and volume when compared to WT counterparts (**Fig. 4C)**, consistent with the decreased MCV observed in our CBC data (**Fig.2A**). We also calculated that *Emd* KO mice also have a modest, yet statistically significant increase in the ratio of surface area to volume (**Fig. 4C**). Together, these data support that *Emd* KO results in a decrease in MCV and suggests a role for emerin in regulating RBC membrane structure in both sexes.

**Figure 4.**
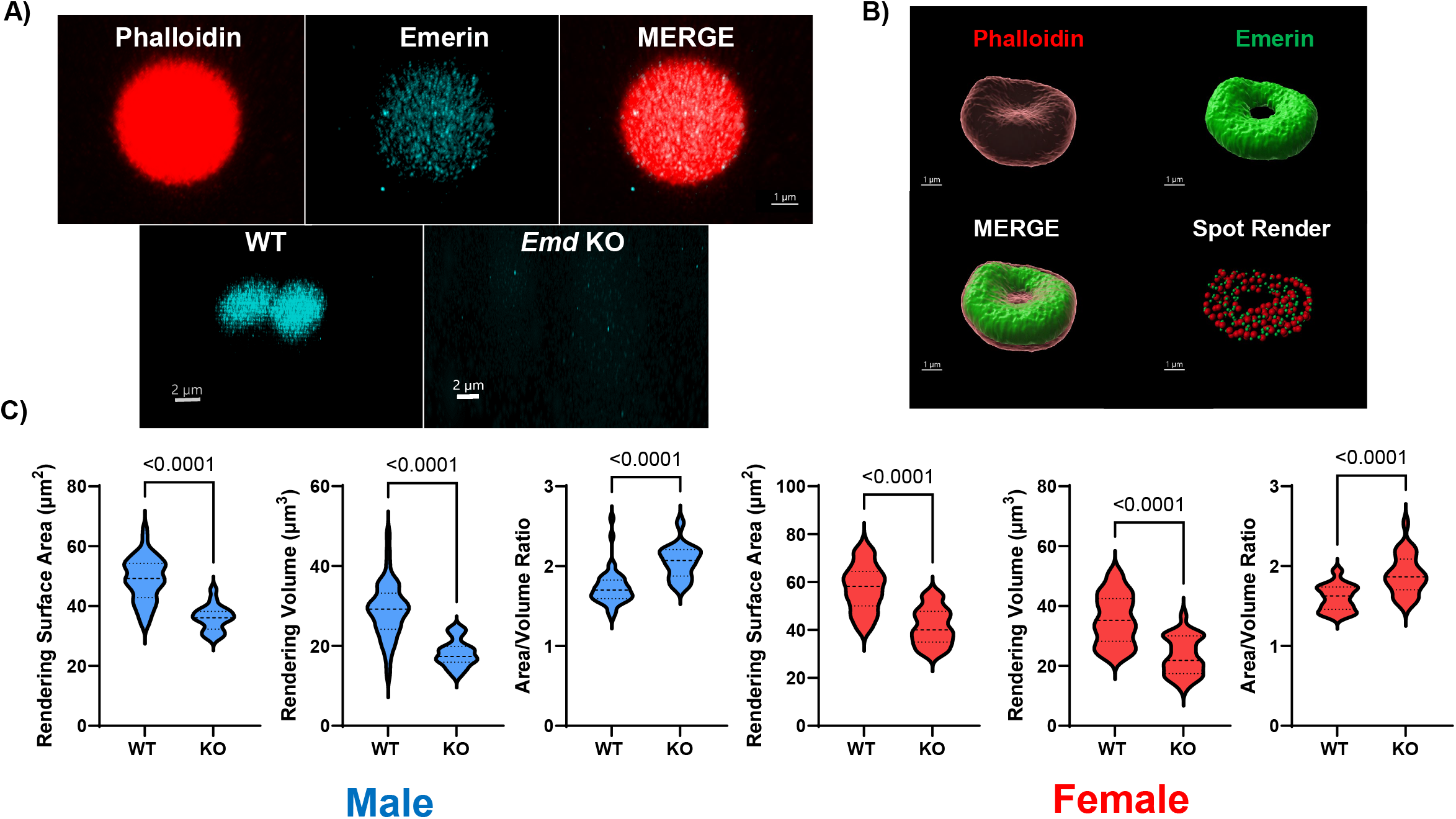
Immunofluorescent labeling and surface rendering to assess emerin-mediated RBC morphology. **A)** Immunofluorescent labeling of RBCs isolated from peripheral blood with AF546-conjugated phalloidin and polyclonal antibody against emerin. Images acquired at 63x with a Leica sp8 inverted confocal microscope. Complete loss of emerin expression was validated by imaging with uniform laser intensity and gain. Scale bar = 1μm (top) and 2μm (bottom). **B)** Imaris surface renderings of z-stack series acquired with the same methodology used in A). Spot render was used to visualize proximity of emerin to phalloidin+ fluorescent loci. **C)** Quantitation of spatial parameters from surface renderings of RBC membrane based on phalloidin staining intensity. Surface area, volume, and calculated A/V ratio were used as a readout of size and shape. Comparisons were made between male (left set, blue) and female (right set, red). Data represented as Mean ± SEM, Student’s t-test.

### Loss of emerin expression does not significantly alter the proportion of erythroid progenitors

To test whether the RBC phenotype observed in the *Emd* KO mice is intrinsic to mature RBCs or is due to a canonical nuclear function in erythroid precursor populations, we isolated hematopoietic progenitor cells from WT and *Emd* KO bone marrow and quantified the percentage of lineage negative cells by FACS (see Methods). We did not detect a significant difference in the percentage of pre-CFU-E or CFU-E cells within lineage negative cells derived from *Emd* KO mice and WT controls. In detection of other hematopoietic progenitors, we found that *Emd* KO resulted in decreased populations of cell types related to the granulocyte and monocyte lineage such as MPP3 and pre-GM along with a decreased percentage of common lymphoid progenitor cells (CLP) (**Fig. 5**).

**Figure 5.**
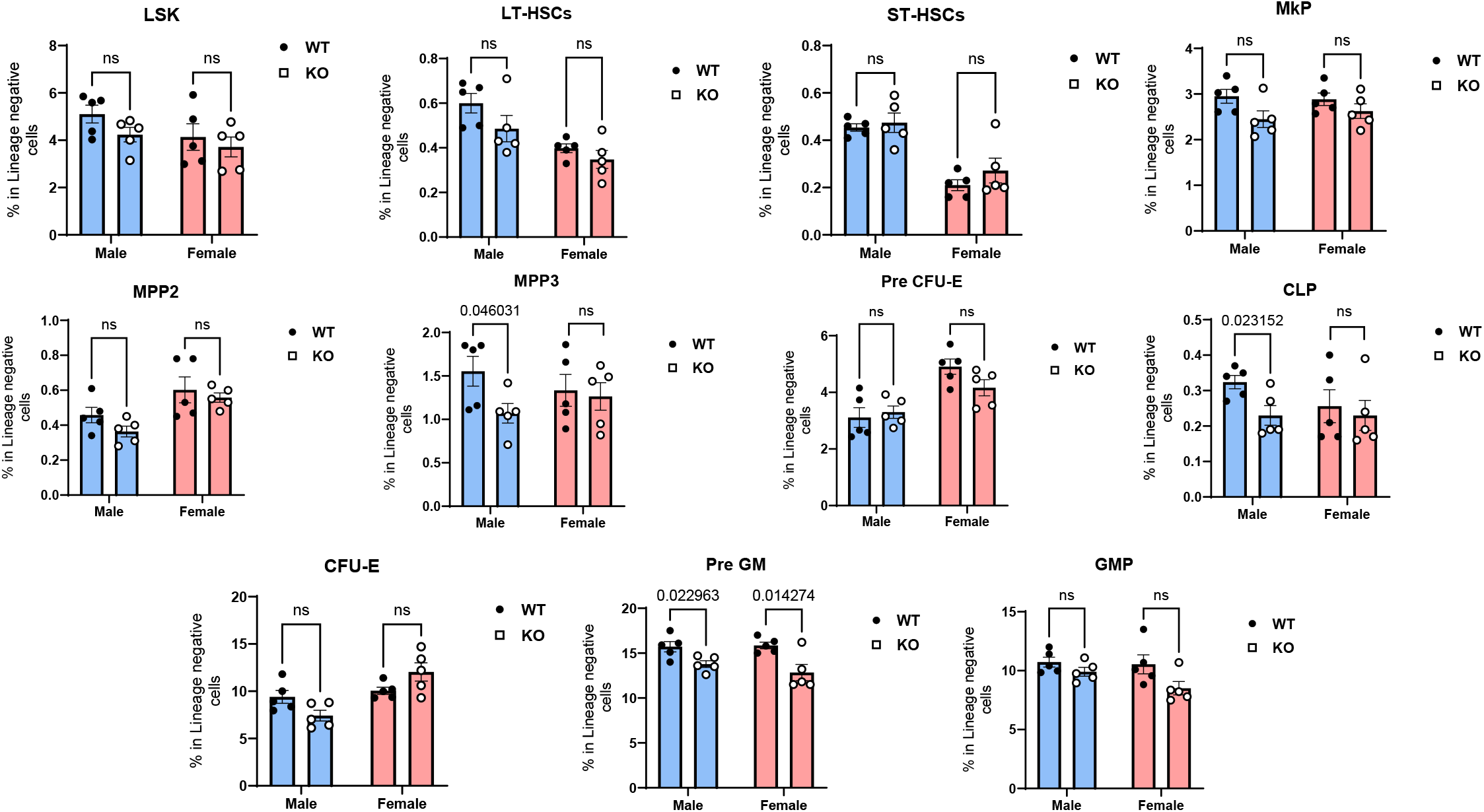
Analysis of lineage-negative hematopoietic cell populations in WT and *Emd*-deficient bone marrow. FACS sorting of hematopoietic stem cells derived from both sexes of WT and *Emd* KO bone marrow. Male mice shown as blue bars, female mice shown as red bars. Antibody-based identification provided in Methods. n = 5 for each sex of each genotype. Data presented as Mean ± SEM. Student’s t-test for genotype comparisons with each sex.

It has been shown in previous publications that emerin is detected at the transcript (**Fig. 6A**) and protein (**Fig. 6B**) levels in differentiating erythroid progenitor cell populations^38,39^. Emerin expression has been shown to decrease during differentiation across progenitor cell types such as G1ER and MEDEP (**Fig. 6B**). We tested whether emerin plays a role in differentiation of the erythroid lineage and identified that *Emd* KO male mice had decreased percentages of proerythroblasts (ProE) and basophilic erythroblast (Baso) relative to WT bone marrow (**Fig. 6C**). We did not observe a significant phenotype in female mice. Taken together, these data suggest that emerin may play a role in proerythroid differentiation but does not have a significant role in commitment to the erythroid lineage, as detected by lack of phenotype related to CFU-E and pre CFU-E populations. Additionally, these data suggest that emerin also plays a significant role in granulocyte differentiation, specifically neutrophils, which is consistent with the CBC data supporting a male-only decrease in circulating neutrophils (**Fig. 3B**).

**Figure 6.**
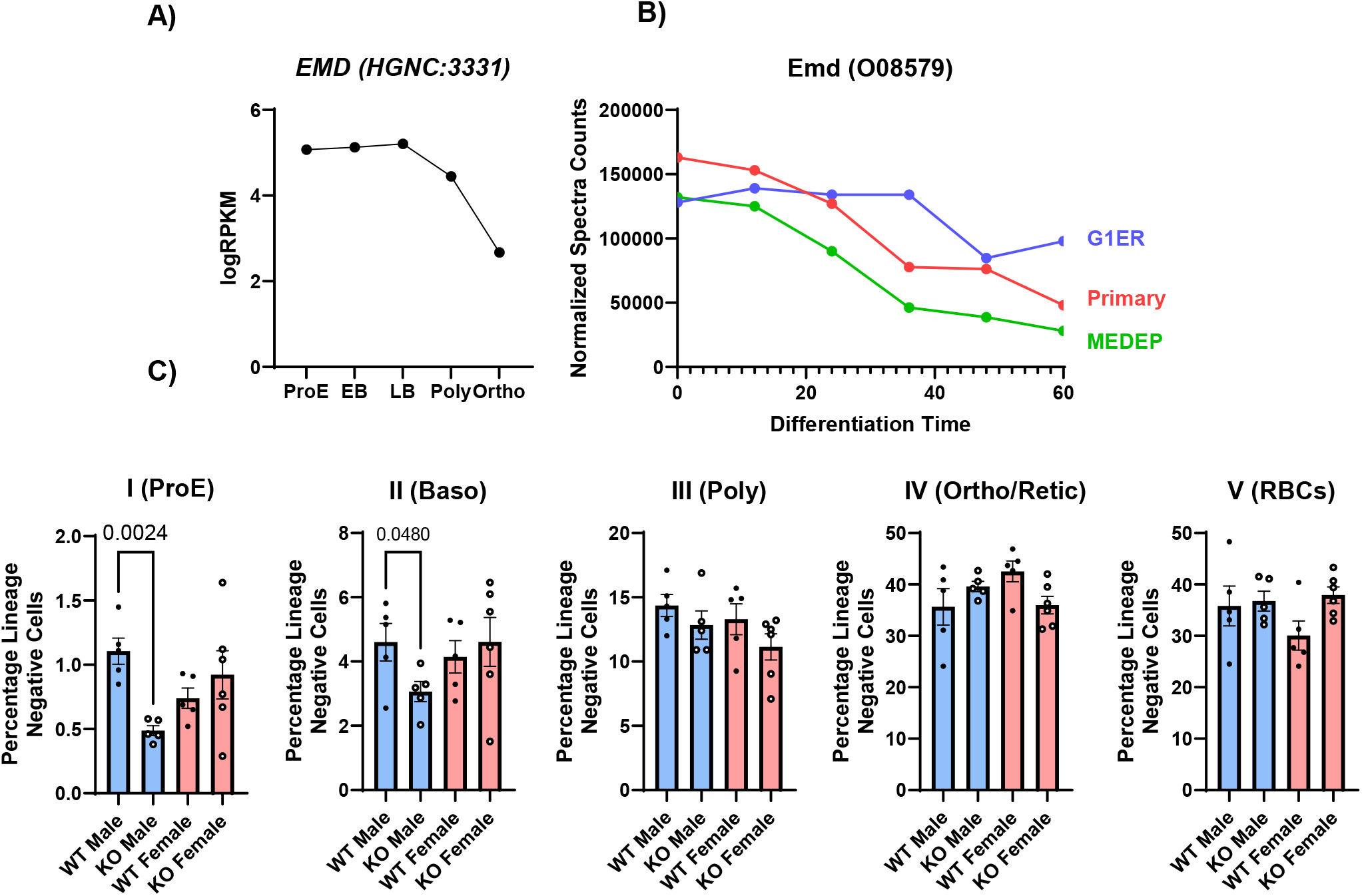
Analysis of erythroid progenitor cell populations in WT and *emerin*-deficient bone marrow. **A)** *EMD* gene expression derived from RNAseq data of erythroid progenitor populations (Gallagher, 2014)^38^. **B)** Proteomic quantitation of EMD peptide abundance during differentiation of erythroid progenitor cell types (Gallagher, 2020)^39^. **C)** Bone marrow cells isolated from both sexes of WT and *Emd* KO mice were stained with an erythroid-focused antibody panel with DAPI for live/dead discrimination (described fully in Methods). Data were quantified as percentage of lineage negative cells in each stage of differentiation. n = 5 for each sex of each genotype. Male mice shown as blue bars, female mice shown as red bars. Data presented as Mean ± SEM. Student’s t-test.

### Emerin-deficient RBCs are more resistant to osmotic hemolysis

Based on our CBC data and microscopy-based quantitation of volume and area, we hypothesized that *Emd* KO cells with decreased MCV would be protected from hemolysis^9^. To test the functional role of emerin in RBC stability and membrane integrity, we performed an osmotic fragility test by exposing RBCs to a series of hypotonic saline solutions. We observed that RBCs from *Emd* KO mice are more resistant to osmotic hemolysis, particularly at 0.55% and 0.6% saline relative to WT counterparts. We observe a more robust phenotype in male mice compared to females (**Fig. 7A**), consistent with the CBC data. We also found that male *Emd* KO mice have a decreased level of free plasma hemoglobin, whereas no significant difference was observed in females relative to their control counterparts (**Fig. 7B**). To test whether *Emd*-dependent RBC membrane integrity is specific to osmotic stress or represents a broader conserved function, we used phenylhydrazine injection as a model of oxidant-mediated hemolysis^40,41^. Under these conditions, we find that *Emd* KO did not have a protective effect on hemolysis following treatment with phenylhydrazine (**Fig. 7C**). Together, these data support our other data that emerin is critical for maintenance of RBC membrane deformity and stability, which protects from osmotic hemolysis, consistent with the observed decrease in MCV.

**Figure 7.**
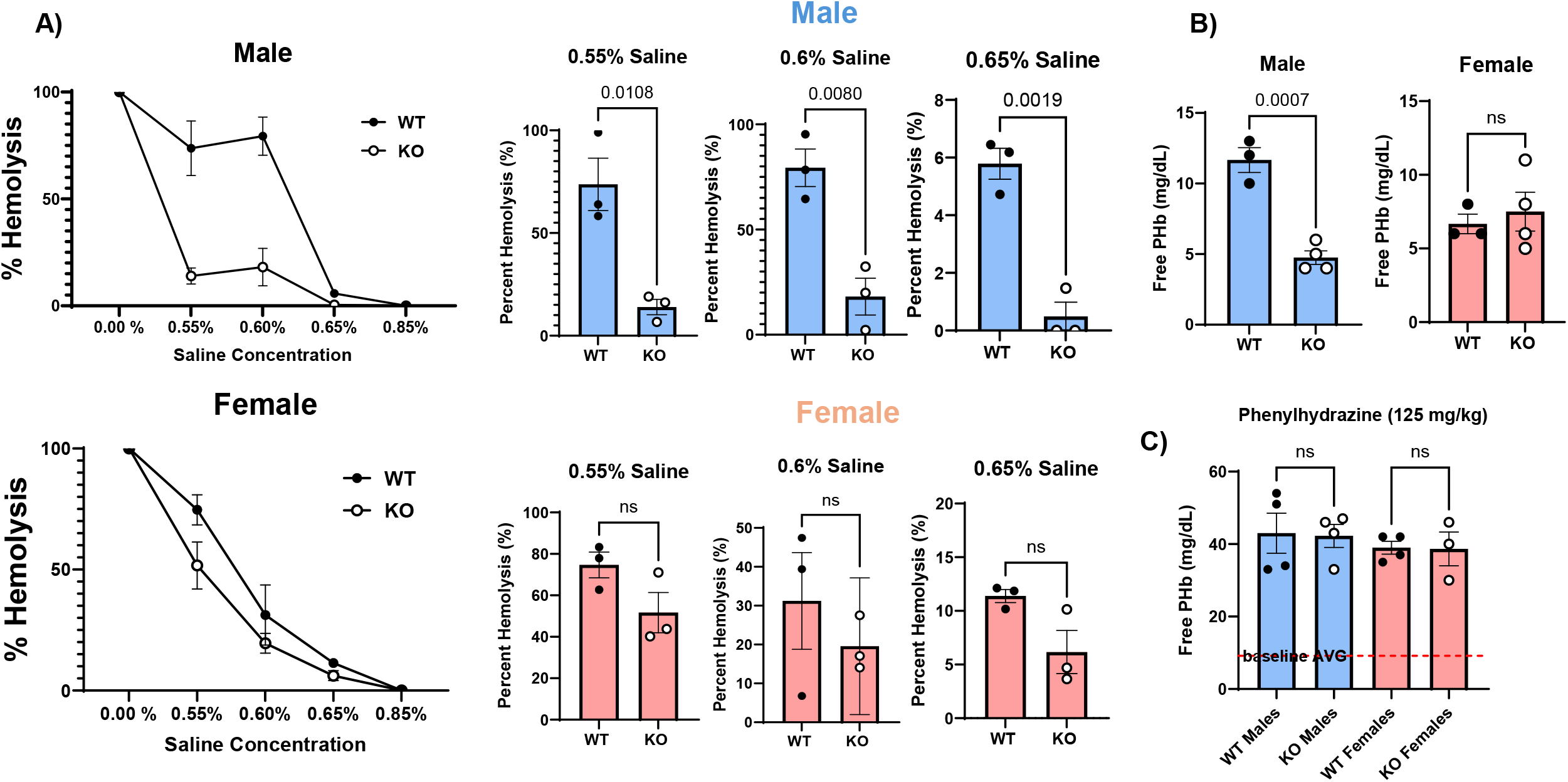
Emerin-deficient RBCs are functionally protected from osmotic hemolysis. **A)** Osmotic gradient fragility assay of peripheral blood RBCs collected from male (top row) and female (bottom row) WT and *Emd* KO mice. Quantitation of hemolysis provided in Methods. n = 3 for each sex of each genotype. Male mice shown as blue bars, female mice shown as red bars. Data presented as Mean ± SEM. Student’s t-test comparing genotypes within sexes at specified saline concentrations. **B)** Free plasma hemoglobin concentration is quantified from isolated plasma. n = 3 WT animals of both sexes, 4 KO animals of both sexes. Data presented as Mean ± SEM. Student’s t-test. **C)** Mice were injected with 125 mg/kg phenylhydrazine 6 hours before blood collection and plasma isolation. Free plasma hemoglobin was quantified and graphed as Mean ± SEM. n = 4 WT males, KO males, and WT females. n = 3 KO females. Data presented as Mean ± SEM. Student’s t-test.

### Human *EMD* variants correlate with altered RBC phenotypes

We next asked whether the RBC phenotype identified in our *Emd* KO mouse model (MCV, MCH) are also altered in individuals with mutations in *EMD*. First, we interrogated the Common Metabolic Disease Knowledge Portal (CMDKP), which aggregates, analyzes, and displays human genetic and functional genomic information linked to common metabolic diseases and traits^42^. We found that human *EMD* mutations have very strong human genetic evidence (HuGE) scores (calculated by common and rare variants) related to mean sphered volume and reticulocyte count and moderate scores related to most CBC parameters (**Fig. 8A**). Next, we employed a machine learning model called Deep Phenotype-Variant Prioritization (Deep PVP) to integrate phenotype data from the ExPheWAS and Pan-UK Biobank and observe hematological phenotypes associated with *EMD* variants. We found that a significant portion of tissue/cell types affected by *EMD* mutations were related to immune/blood cells and that there are statistically significant effects on RBC indices, including mean sphered cell volume, reticulocyte parameters, and mean corpuscular volume (**Fig. 8B**). Finally, we asked whether individuals clinically diagnosed with X-linked Emery-Dreifuss muscular dystrophy (EDMD) also have RBC phenotypes consistent with those observed in our mouse model. CBC data was collected and analyzed from five male patients diagnosed with EDMD validated by loss of emerin expression (IF and/or western blot, performed on lymphoid cell lines or muscle biopsies). These data highlight a phenotype that is consistent with our previous findings, most notably with a modest, yet reproducible decrease in MCV. The values fall within the normal reference ranges (red dashed lines) in adult males and are approximately one standard deviation from the mean (**Figure 8C**) potentially explaining why none of these patients had RBC shape abnormalities or clinical symptoms such anemia. Overall, these data are consistent with our *in vivo* data finding that emerin plays a novel non-nuclear role in regulating plasma membrane structure and integrity in RBCs and is a previously unidentified modifier of RBC traits.

**Figure 8.**
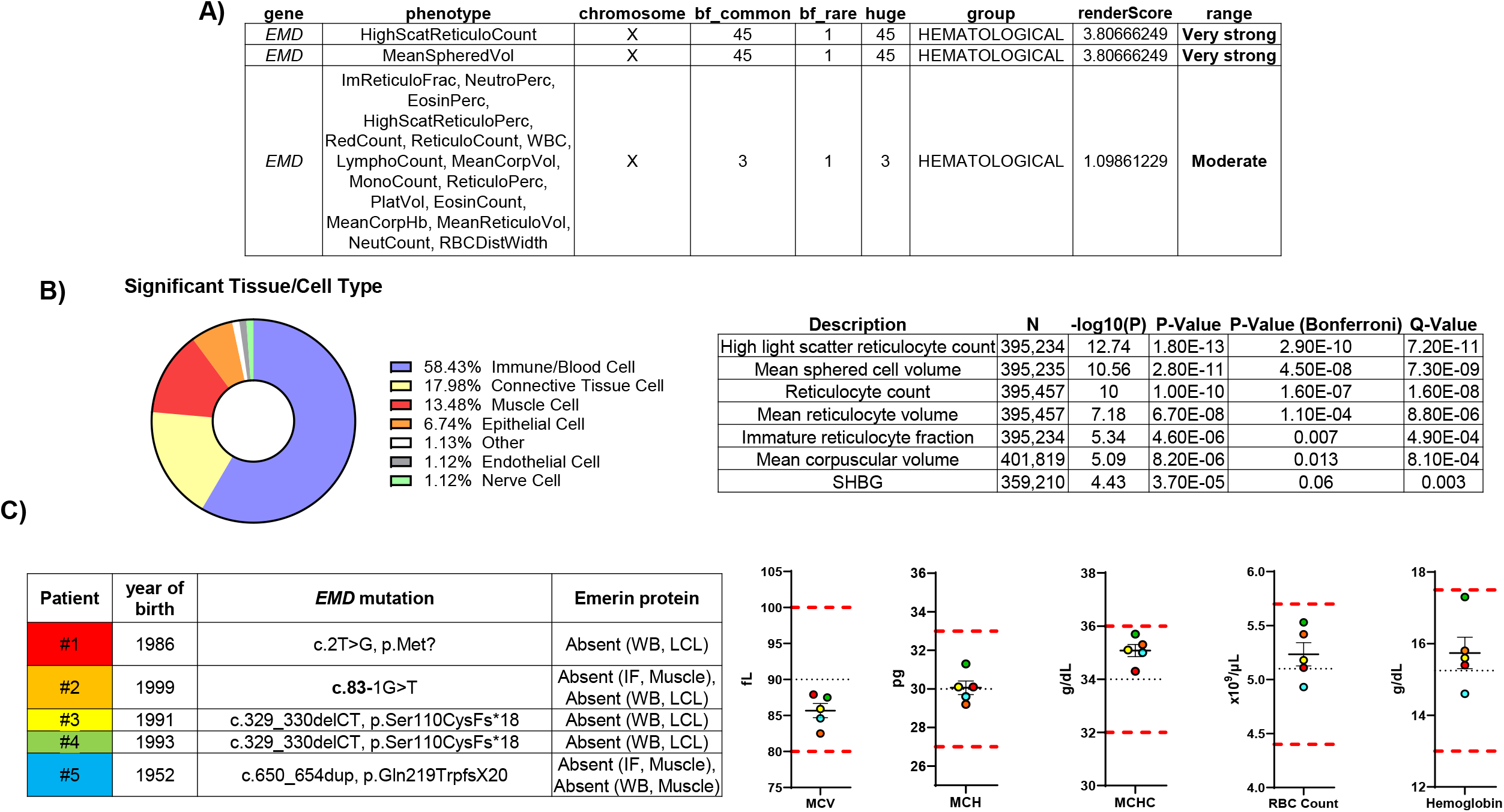
Human *EMD* mutations are associated with RBC phenotypes. **A)** Red cell/hematological enrichment associated with human *EMD* mutations according to query of the Common Metabolic Disease Knowledge Portal (CMDKP) represented as their calculated human genetic evidence (HuGE) score. **B)** Deep Phenotype-Variant Prioritization (Deep PVP) was utilized to extract genotypic and phenotypic associations from the ExPheWAS platform and the Pan-UK Biobank. Significantly enriched cell types (left) and enrichment of RBC-related parameters (right) are displayed. **C)** CBC data from individuals diagnosed with X-linked Emery-Dreifuss muscular dystrophy (EDMD). Individual patient statistics and validation of emerin deficiency are provided (left) and color-coded. RBC parameters (right) are compared to physiological human ranges (red dashed lines) and average values (black dashed lines). IF: immunofluorescence analysis, WB: western blot emerin analysis, LCL: lymphoblastoid cell lines.

## Discussion

Our study reveals a previously unidentified role for emerin, a well-characterized nuclear envelope protein, in regulating the structure and function of mature red blood cells (RBCs). The data presented here support that emerin is a major component of the mature RBC membrane with functional roles in cytoskeletal organization, RBC morphology, and resistance to osmotic stress.

We first demonstrated that emerin is expressed in mature, anucleate RBCs and enriched in the plasma membrane, suggesting that its function extends beyond its canonical roles regulating nuclear architecture and gene expression. Proteomic analyses revealed significant decrease in expression of protein complexes involved in actin dynamics, septin assembly, and cytoskeletal organizations in emerin-deficient mice, providing molecular evidence that emerin contributes to membrane structural integrity by associating with cytoskeletal networks. These associations are especially compelling given the critical role of the cytoskeleton in maintaining RBC deformability and survival in circulation, which were further validated by the phenotype ontology analysis (Monarch) highlighting RBC membrane dysfunction as a pathway affected by emerin expression.

From these data, we tested the functional consequence of *Emd* deletion in our complete blood count (CBC) and imaging analyses. Loss of emerin resulted in reduced MCV and MCH without changes in MCHC, suggesting a smaller but proportionally hemoglobin-rich RBC phenotype. High-resolution microscopy and surface rendering supported the observed decreased cell volume and surface area in *Emd* KO RBCs, supporting a cell-intrinsic structural defect. These data also allowed us to quantify a relative increase in the surface area to volume ratio, suggesting the cells have a more flattened morphology. Since the animals used in this study are full genetic knockouts, this observed MCV phenotype occurred in both male and female mice, despite the X-linked nature of *Emd* gene. Despite this, we did observe a male-only increase in RBC number, hemoglobin, and hematocrit.

Aside from RBCs, our data also highlighted a male-specific reduction in circulating neutrophils in *Emd* KO mice. Given the distinctive nuclear morphology and deformability required for neutrophil function, this result suggests that emerin may also influence nuclear mechanics in other lineages and may contribute to immune cell differentiation. Supporting this, hematopoietic profiling revealed reductions in granulocyte-monocyte progenitor populations (MPP3, pre-GM) and common lymphoid progenitors. While *Emd* deletion did not alter early erythroid progenitor numbers, we observed sex-specific changes in downstream erythroid differentiation stages, including reduced ProE and basophilic erythroblasts in male mice. These findings suggest that emerin may be dispensable for erythroid lineage commitment but modulates late-stage maturation in a sex-dependent manner.

Functionally, emerin-deficient RBCs exhibited decreased susceptibility to osmotic hemolysis, consistent with their reduced MCV and increased surface area-to-volume ratio. This protective effect appears to be specific to osmotic stress, as emerin loss did not confer resistance to oxidant-induced hemolysis *via* phenylhydrazine. These data reinforce the notion that emerin plays a structural role in modulating membrane stability and suggest that its absence leads to smaller, less deformable cells that are more resistant to osmotic rupture.

Leveraging human bioinformatic strategies CMDKP and DeepPVP revealed associations between *EMD* variants and hematologic traits, including mean sphered volume and reticulocyte indices. Interestingly, DeepPVP analysis revealed a statistically significant *EMD* phenotype related to sex hormone binding globulin (SHBG), a circulating protein synthesized in the liver that maintains the balance of sex hormones in the blood, namely testosterone and estradiol^43^. It is possible that the male-dominated phenotypes we observe in our mouse model are partially due to aberrant availability or function of SHBG, which can increase RBC number and hemoglobin independent of the RBC-intrinsic phenotypes observed in both sexes such as MCV. Future investigations are ongoing to differentiate endocrine and hematopoietic contributions to the observed *EMD* phenotype. Although EDMD is a rare disease with an estimated prevalence of 1.3:100,000-2:100,000^44,45^, we were able to retroactively analyze the CBC data from a small cohort of male patients with confirmed X-linked EDMD and found that they exhibited mild but consistent reductions in MCV, recapitulating the phenotype observed in *Emd* KO mice. While these changes fall within normal reference ranges, they suggest that *EMD* gene is a previously underappreciated genetic modifier of red cell traits in humans.

Overall, this work identifies a novel, extranuclear role for emerin in maintaining RBC membrane architecture and mechanical stability. Our findings broaden the biological scope of emerin beyond its established nuclear roles and implicate it in the regulation of cytoskeletal integrity in anucleate cells. These results prompt reconsideration of RBC traits in patients with *EMD* mutations and suggest new avenues for investigating hematologic pathobiology in X-linked EDMD and related disorders. Future studies are be needed to determine how the molecular mechanisms of emerin-cytoskeleton interaction govern RBC morphology and to explore the broader hematopoietic implications of emerin deficiency, including its role in immune cell differentiation and function.

## Acknowledgements

We would like to thank the MS & Proteomics Resource at Yale University, namely Katie Henke, PhD and Jean Kanyo, for providing the necessary mass spectrometers and the accompany biotechnology tools funded in part by the Yale School of Medicine and by the Office of The Director, National Institutes of Health (S10OD02365101A1, S10OD019967, and S10OD018034). The funders had no role in study design, data collection and analysis, decision to publish, or preparation of the manuscript. Additionally, we would like to acknowledge Rolando Garcia Milian MLS, AHIP at the Yale Medical Library for his bioinformatic expertise. We also would like to thank the veterinary staff at the Yale Animal Resource Center for their commitment to the health and wellbeing of the animals used in this investigation. We finally thank the clinicians, Dr Tanya Stojkovic, Dr Pascal Laforêt, who follow *EMD* mutated patients, for providing CBC data.

## Authorship Contributions

Brendan O’Brien designed and performed most experiments listed, wrote, and edited the manuscript. Sean Gu served as an expert consultant on experimental design and performed all the blood draws and coordination with Yale Laboratory Medicine. Kathleen Martin, and John Hwa provided reagents and expert insight regarding experimental design, interpretation of results, and editing the manuscript. Timothy Nottoli and Suxia Bai designed and created the CRISPR-edited mouse strain. Patrick Gallagher and Joyleen Oliver served as expert consultants in RBC physiology/membrane disorders and provided significant editing to the manuscript. Evrett Thompson, Madeline Mayday, Rubia Mancuso, and Diane Krause helped perform the bone marrow hematopoietic stem cell FACS and analysis. Gisèle Bonne, Rabah Ben Yaou, and France Leturcq provided expert insight into emerin/nuclear envelope biology and critical patient data from males diagnosed with X-linked-EDMD. Jennifer Kwan and Alokkumar Jha provided bioinformatics analysis of UK Biobank data regarding RBC phenotypes in individuals with mutations in *EMD*.

